# A Genealogical Look at Shared Ancestry on the X Chromosome

**DOI:** 10.1101/046912

**Authors:** Vince Buffalo, Stephen M. Mount, Graham Coop

## Abstract

Close relatives can share large segments of their genome identical by descent (IBD) that can be identified in genome-wide polymorphism datasets. There are a range of methods to use these IBD segments to identify relatives and estimate their relationship. These methods have focused on sharing on the autosomes, as they provide a rich source of information about genealogical relationships. We can hope to learn additional information about recent ancestry through shared IBD segments on the X chromosome, but currently lack the theoretical framework to use this information fully. Here, we fill this gap by developing probability distributions for the number and length of X chromosome segments shared IBD between an individual and an ancestor *k* generations back, as well as between half-and full-cousin relationships. Due to the inheritance pattern of the X and the fact that X homologous recombination only occurs in females (outside of the pseudo-autosomal regions), the number of females along a genealogical lineage is a key quantity for understanding the number and length of the IBD segments shared amongst relatives. When inferring relationships among individuals, the number of female ancestors along a genealogical lineage will often be unknown. Therefore, our IBD segment length and number distributions marginalize over this unknown number of recombinational meioses through a distribution of recombinational meioses we derive. We show how our results can be used to estimate the number of female ancestors between two relatives, giving us more genealogical details than possible with autosomal data alone.

---

Close relatives are expected to share large contiguous segments of their genome due to the limited number of crossovers per chromosome each generation (Donnelly, 1983; Fisher, 1949, 1954). These large identical by descent (IBD) segments shared among close relatives leave a conspicuous footprint in population genomic data, and identifying and understanding this sharing is key to many applications in biology (Thompson, 2013). For example, in human genetics, evidence of recent shared ancestry is an integral part of detecting cryptic relatedness in genome-wide association studies (Gusev et al., 2009), discovering mis-specified relationships in pedigrees (Sun et al., 2002), inferring pairwise relationships (Glaubitz et al., 2003; Huff et al., 2011), and localizing disease traits in pedigrees (Thomas et al., 2008). In forensics, recent ancestry is crucial for both accounting for population-level relatedness (Balding et al., 1994) and in familial DNA database searches (Belin et al., 1997; Sjerps et al., 1999). Additionally, recent ancestry detection methods have a range of applications in anthropology and ancient DNA to understand the familial relationships among sets of individuals (Baca et al., 2012; Fu et al., 2015; Haak et al., 2008; Keyser-Tracqui et al., 2003). In population genomics, recent ancestry has been used to learn about recent migrations and other demographic events (Palamara et al., 2012; Ralph et al., 2013). An understanding of recent ancestry also plays a large role in understanding recently admixed populations, where individuals draw ancestry from multiple distinct populations (Gravel, 2012; Liang et al., 2014; Pool et al., 2009). Finally, relative finding through recent ancestry is increasingly a key feature of direct-to-consumer personal genomics products and an important source of information for genealogists leveraging genetic information (Durand et al., 2014; Royal et al., 2010).

Approaches to infer recent ancestry among humans have often used only the autosomes, as the recombining autosomes offer more opportunity to detect a range of relationships than the Y chromosome, mitochondria, or X chromosome. However, the nature of X chromosome inheritance means that it can clarify details of the relationships among individuals and be informative about sex-specific demography and admixture histories in ways that autosomes cannot (Bryc et al., 2010; Bustamante et al., 2009; Goldberg et al., 2015; Pool et al., 2007; Ramachandran et al., 2008, 2004; Shringarpure et al., 2016).

In this paper, we look at the inheritance of chromosomal segments on the X chromosome among closely related individuals. Our genetic ancestry models are structured around biparental genealogies back in time, an approach used by many previous authors (e.g., Barton et al., 2011; Chang, 1999; Donnelly, 1983; Rohde et al., 2004). If we ignore pedigree collapse (due to inbreeding), the genealogy of a present-day individual encodes all biparental relationships back in time; e.g. the two parents, four grandparents, eight great-grandparents, 2^*k*^ great^*k*-2^ grandparents, and in general the 2^*k*^ ancestors *k* generations back; we refer to these individuals as one’s *genealogical ancestors*. A genealogical ancestor of a present-day individual is said to also be a *genetic ancestor* if the present-day individual shares genetic material by descent from this ancestor. We refer to these segments of shared genetic material as being identical by descent, and in doing so we ignore the possibility of mutation in the limited number of generations separating our individuals. Throughout this paper, we will ignore the pseudo-autosomal (PAR) region of the X chromosome, which undergoes crossing over with the Y chromosome in males (Koller et al., 1934) to ensure proper disjunction in meiosis I (Hassold et al., 1991). We also ignore gene conversion which is known to occur on the X (Rosser et al., 2009).

Here, we are concerned with inheritance through the *X genealogy* embedded inside an individual’s genealogy, which includes only the subset of one’s genealogical ancestors who could possibly contributed to one’s non-PAR X chromosome. We refer to the individuals in this X genealogy as *X ancestors*. Since males receive an X only from their mothers, a male’s father cannot be an X ancestor. Consequently, a male’s father and all of his ancestors are excluded from the X genealogy (Figure 1). Therefore, females are overrepresented in the X genealogy, and as we go back in one’s genealogy, the fraction of individuals who are possible X ancestors shrink. This property means that genetic relationships differ on the X compared to the autosomes, a fact that changes the calculation of kinship coefficients on the X (Pinto et al., 2011, 2012) and also has interesting implications for kin-selection models involving the X chromosome (Fox et al., 2009; Rice et al., 2008).

**Figure 1:**
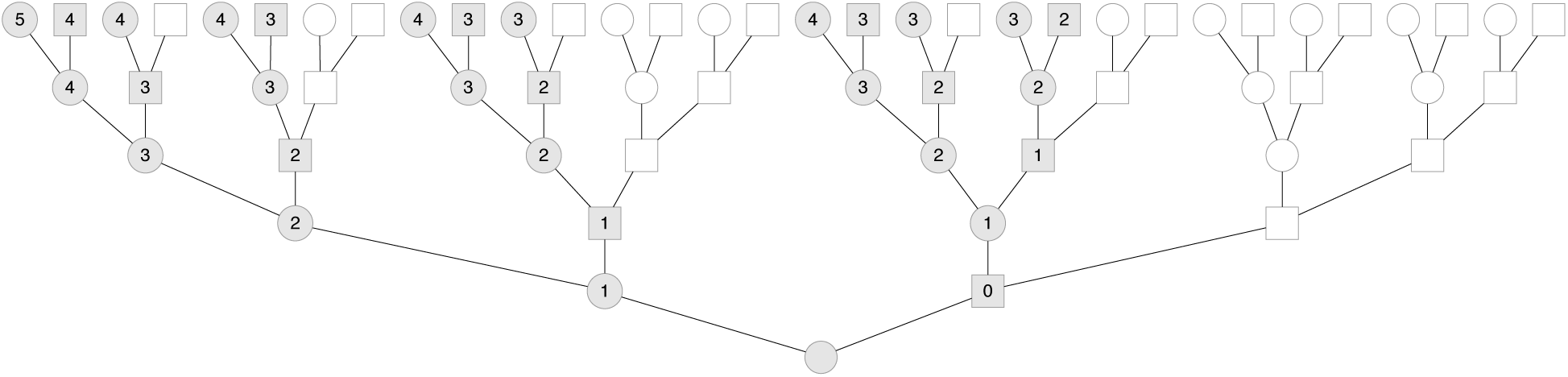
An example genealogy with the embedded X genealogy back five generations. Males are depicted as squares and females as circles. Individuals in the X genealogy are shaded gray while unshaded individuals are ancestors that are not X ancestors. Each X ancestor is labeled with the number of recombinational meioses to the present-day female.

In Section 1 we review models of autosomal identity by descent among relatives, on which we base our models of X genetic ancestry. Then, in Section 2 we look at X genealogies back in time, as the properties of X genealogies affect the transmission of X genetic material from X ancestors to a present-day individual. We develop simple approximations to the probability distributions of the number and length of X chromosome segments that will be shared IBD between a present-day female and one of her X ancestors a known number of generations back in time. These models provide a set of results for the X chromosome equivalent to those already known for the autosomes (Donnelly, 1983; Thomas et al., 1994). Then, in Section 3, we look at shared X ancestry—when two present-day cousins share an X ancestor a known number of generations back. We calculate the probabilities that genealogical half-and full-cousins are also connected through their X genealogy, and thus can potentially share genetic material on their X. We then extend our models of IBD segment length and number to segments shared between half-and full-cousins. Finally, in Section 4 we show that shared X genetic ancestry contains additional information (compared to genetic autosomal ancestry) for inferring relationships among individuals, and explore the limits of this information.

## 1 Autosomal Ancestry

To facilitate comparison with our X chromosome results, we first briefly review analogous autosomal segment number and segment length distributions (Donnelly, 1983; Huff et al., 2011; Thomas et al., 1994). Throughout this paper, we assume that one’s genealogical ancestors *k* generations back are distinct, i.e. there is no pedigree collapse due to inbreeding (see Appendix D for a model of how this assumption breaks down with increasing *k*). Thus, an individual has 2^*k*^ *distinct* genealogical ancestors. Assuming no selection and fair meiosis, a present-day individual’s autosomal genetic material is spread across these 2^*k*^ ancestors with equal probability, having been transmitted to the present-day individual solely through recombination and fair segregation.

We model the process of crossing over during meiosis as a continuous time Markov process along the chromosome, as in Thomas et al. (1994) and Huff et al. (2011), and described by Donnelly (1983). In doing so we assume no crossover interference, such that in each generation *b* recombinational breakpoints occur as a Poisson process running with a uniform rate equal to the total length of the genetic map (in Morgans), *v.* Within a single chromosome, *b* breaks creates a mosaic of *b* + 1 alternating pieces of maternal and paternal segments. This alternation between maternal and paternal haplotypes creates long-run dependency between segments. We ignore these dependencies in our analytic models by assuming that each chromosomal segment survives segregation independently with probability ½ per generation. For *d* independent meioses separating two individuals, we imagine the Poisson recombination process running at rate *vd*, and for a segment to be shared IBD between the two ancestors it must survive 1/2^*d*^ segregations. Consequently, the expected number of segments shared IBD between two individuals *d* meioses apart in a genome with *c* chromosomes is approximated as (Thomas et al., 1994):

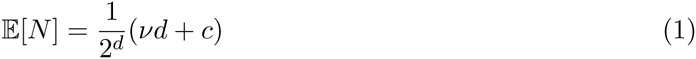

Intuitively, we can understand the 1*/* 2*d* factor as the coefficient of kinship (or path coefficient; Wright, 1922, 1934) of two individuals *d* meioses apart, which gives the probability that two alleles are shared IBD between these two individuals. Then, the expected number of IBD segments 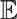[*N*] can be thought of as the average number of alleles shared between two individuals in a genome with *vd* + *c* loci total. Under this approximation, recombination increases the number of independent loci linearly each generation (by a factor of the total genetic length). A fraction 1/2^*d*^ of parental alleles at these loci survive the *d* meioses to be IBD with the present-day individual.

By convention, we count the number of contiguous IBD segments *N* in the present-day individual, not the number of contiguous segments in the ancestor. For example, an individual will share exactly one block per chromosome with each parent if we count the contiguous segments in the offspring, even though these segments may be spread across the parent’s two homologues. This convention, which we use throughout the paper, is identical to counting the number of IBD segments that occur in *d*-1 meioses rather than *d* meioses. This convention only impacts models of segments shared IBD between an individual and one of their ancestors; neither the distribution of segment lengths nor the distributions for segment number shared IBD between cousins are affected by this convention.

### The distribution of IBD segments between a present-day individual and an ancestor

Given that a present-day individual and an ancestor in the *k*^th^ generation are separated by *k* meioses, the number of IBD segments can be modeled with what we call the *Poisson-Binomial* approximation. Over *d* = *k* meioses, *B* = *b* ∼ Pois(*vk*) recombinational breakpoints fall on *c* independently assorting chromosomes, creating *b* + *c* segments. Ignoring long-range dependencies, we assume all of these *b* + *c* segments have an independent chance of surviving the *k* segregations to the present-day individual, and thus the probability that *n* segments survive given *b* + *c* trials is Binomially distributed with probability 1/2^*k*^. Marginalizing over the unobserved number of recombinational breakpoints *b*, and replacing *k* with *k* - 1 to following the convention described above:

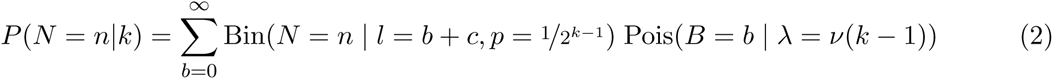

The expected value of the Poisson-Binomial model is given by equation (1) and this model is similar to those of Donnelly (1983) and Thomas et al. (1994). We can further approximate this by assuming that we have a Poisson total number of segments with mean (*c* + *v*(*k* - 1)) and these segments are shared with probability 1/2^*k-1*^ as in Huff et al. (2011). This gives us a thinned Poisson distribution of shared segments:

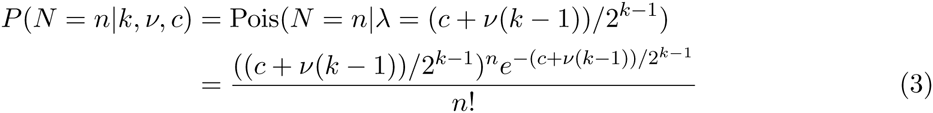

This thinned Poisson model also has an expectation given by equation (1) but compared to the Poisson-Binomial model has a larger variance than the true process. This overdispersion occurs because modeling the number of segments created after *b* breakpoints involves incorporating the initial number of chromosomes into the Poisson rate. However, this initial number of chromosomes is actually fixed, which the Poisson-Binomial model captures but the Poisson thinning model does not (i.e. one generation back such that *k* = 1, the thinning model treats the number of segments shared IBD with one’s parents is *N* ∼ Pois(*c*) rather than *c*). See Appendix A for a further comparison of these two models. A more formal description of this approximation as a continuous-time Markov process is given in Thomas et al. (1994).

### The distribution of IBD segments between cousins

Similarly, we can derive the distribution for the number of IBD segments shared between two half-cousins with an ancestor in the *k*^th^ generation. Two half-cousins are separated by 2*k* meioses, thus the distribution for number of segments is:

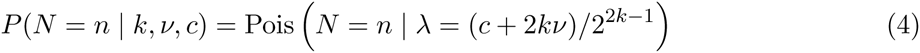

Since either of the shared parent’s haplotypes can be shared IBD between the two cousins, the Poisson process rate is doubled. Full-cousins can share segments via either of their two shared ancestors, leading the distribution to be:

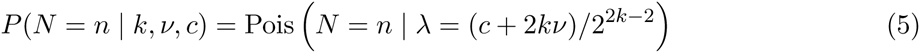

### The distribution of autosome segment lengths

In addition to the number of IBD segments, the length of segments is also informative about ancestry (e.g. Palamara et al., 2012). As we model crossing over as a Poisson process, a one Morgan region will experience on average *d* recombination events over *d* meioses. Therefore, the probability density of segment lengths shared IBD between two individuals *d* meioses apart is exponential with rate *d*:

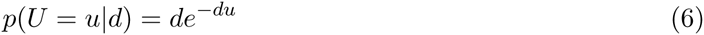

Equations (6) and (4) specify a model of the number and lengths of segments shared between various degree relatives. Various authors have used these types of results to derive likelihood-based models for classifying the genealogical relationship between pairs of individuals using autosome IBD data (Durand et al., 2014; Henn et al., 2012; Huff et al., 2011).

We will use similar models as these in modeling the length and number of X chromosome segments shared been relatives. However, the nature of X genealogies (which we cover in the next section) requires we adjust these models. Specifically, while one always has *k* recombinational meioses between an autosomal ancestor in the *k*^th^ generation, the number of recombinational meioses varies across the lineages to an X ancestor with the number of females in a lineage, since X homologous recombination only occurs in females (Figure 1). This varying number of recombinational meioses across lineages leads to a varying-rate Poisson recombination process, with the rate depending on the specific lineage to the X ancestor. After we take a closer look at X genealogies in the next section, we adapt the models above to handle the varying-rate Poisson process needed to model IBD segments in X genealogies.

## 2 X Ancestry

### Number of Genealogical X Ancestors

While a present-day individual can potentially inherit autosomal segments from any of its 2^*k*^ genealogical ancestors *k* generations back, only a fraction of these individuals can possibly share segments on the X chromosome. In contrast to biparental genealogies, males have only one genealogical X ancestor—their mothers—if we ignore the PAR. This constraint (which we refer to through as the *no two adjacent males condition*) shapes both the number of X ancestors and the number of females along an X lineage. For example, consider a present-day female’s X ancestors one generation back: both her father and mother contribute X chromosome material. Two generations back, she has three X genealogical ancestors: her father only inherits an X from her paternal grandmother, while her mother can inherit X material from either of her maternal grandparents. Continuing this process, this individual has five ancestors three generations back and eight ancestors four generations back (Figure 1).

In general, a present-day female’s X genealogical ancestors is growing as a Fibonacci series (Basin, 1963; Laughlin, 1920), such that *k* generations back she has *Ƒ*_*k*+2_ X genealogical ancestors, where *Ƒ*_*k*_ is the *k*^th^ Fibonacci number (OEIS A000045; Sloane, 2010). We can demonstrate that one’s number of X genealogical ancestors grows as a Fibonacci series by encoding the X inheritance rules for the number of males and females (*m*_*k*_ and *f*_*k*_, respectively) in the *k*^th^ generation as a set of recurrence relations:

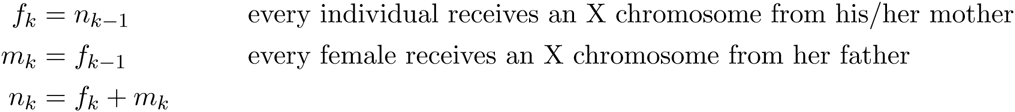

Rearranging these recurrence equations gives us *n*_*k*_ = *n*_*k-1*_ + *n*_*k-2*_, which is the Fibonacci recurrence. Starting with a female in the *k* = 0 generation, we have initial values *n*_0_ = 1 and *n*_1_ = 2, which gives us the Fibonacci numbers offset by two, *Ƒ*_*k*+2_. For a present-day male, his number of X ancestors is *Ƒ*_*k*+1_, i.e. offset by one to count the number of X ancestors his mother has. To simplify our expressions, we will assume throughout the paper that all-present day individuals are female since a simple offset can be made to handle males.

In Figure 3A we show the increase in the number of X genealogical and genetic ancestors and compare these to the growth of all of one’s genealogical ancestors and autosomal genetic ancestors. The closed-form solution for the *k*^th^ Fibonacci number is given by Binet’s formula, which shows that the Fibonacci sequence grows at an exponential rate slower than 2^*k*^.

**Figure 3:**
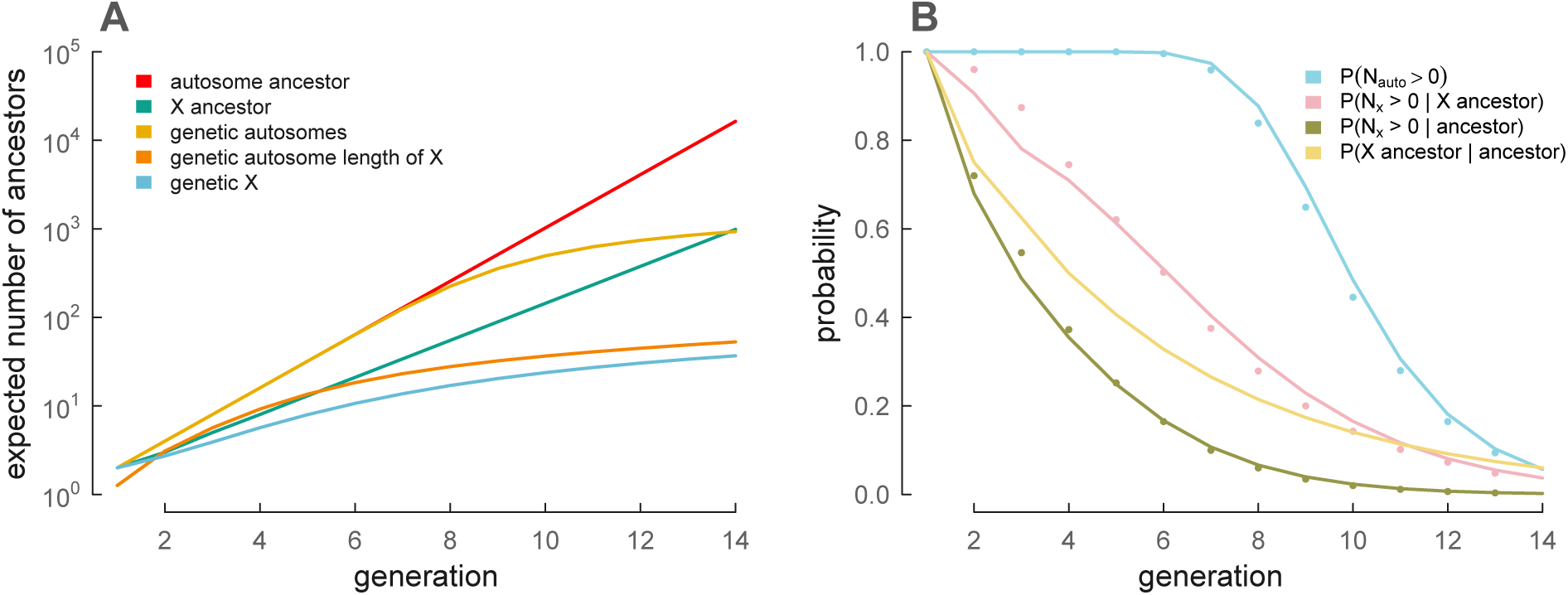
A: A present-day female’s number of genealogical and genetic ancestors, for X chromosomes and autosomes. For comparison to an X chromosome, the number of genetic ancestors of an autosome of length equal to the X is included. B: The probability of genealogical and genetic ancestry for a variety of cases. *P*(N_auto_ > 0) is derived from equation (3), *P*(N_X_ > 0 | X ancestor) from equation (12), *P*(N_X_ > 0 | ancestor) from equations (12) and (7), and *P*(X ancestor | ancestor) from equation (7). Points show simulated results.

Consequently, the fraction of ancestors who can contribute to the X chromosome is declining. Given that a female has *Ƒ*_*k*+2_ X ancestors and 2^*k*^ genealogical distinct ancestors, her proportion of X ancestors is:

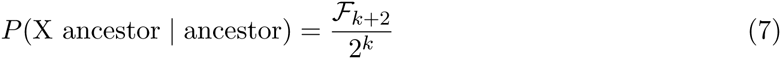

This fraction can also be interpreted as the probability that a randomly chosen genealogical ancestor *k* generations ago is also an X genealogical ancestor. We show this probability as a function of generations into the past in Figure 3B (yellow line).

From our recurrence equations we can see that a present-day female’s *Ƒ*_*k*+2_ ancestors in the *k*^th^ generation are composed of *Ƒ*_*k*+1_ females and *F*_*k*_ males. Likewise for a present-day male, his *Ƒ*_*k*+1_ ancestors in the *k*^th^ generation are composed of *F*_*k*_ females and *Ƒ*_*k*-1_ males. We will use these results when calculating the probability of a shared X ancestor.

### Ancestry Simulations

In the next sections, we use stochastic simulations to verify the analytic approximations we derive; here we briefly describe the simulation methods. We have written a C and Python X genealogy simulation procedure (source code available in the supplement and at https://github.com/vsbuffalo/x-ancestry/). We simulate a female’s X chromosome genetic ancestry back through her X genealogy. Each simulation begins with two present-day female X chromosomes, one of which is passed to her mother and one to her father. Segments transmitted to a male ancestor are simply passed directly back to his mother (without recombination). For segments passed to a female ancestor, we place a Poisson number of recombination breakpoints (with mean *v*) on the X chromosome and the segment is broken where it overlaps these recombination events. The first segment along the chromosome is randomly drawn to have been inherited from either her mother/father, and we alternate this choice for subsequent segments. This procedure repeats until the target generation back *k* is reached. The segments in the *k*-generation ancestors are then summarized as either counts (number of IBD segments per individual) or lengths. These simulations are necessarily approximate as they ignore crossover interference. However, unlike our analytic approximations, our simulation procedure maintains long-run dependencies created during recombination, allowing us to see the extent to which assuming independent segment survival adversely impacts our analytic results.

Figure 2A depicts the X genetic ancestors of one simulated example X genealogy back nine generations to illustrate this process. Each arc represents a single ancestor of a present-day female; red arcs are female X ancestors and blue arcs are male X ancestors. Ancestors that are not X ancestors are shaded gray. The saturation indicates the genetic contribution of this individual to the present-day female’s X (with white indicating an X ancestor that made no genetic contribution on the X). Figure 2B shows the underlying X chromosome IBD segments for this same simulation back four generations.

**Figure 2:**
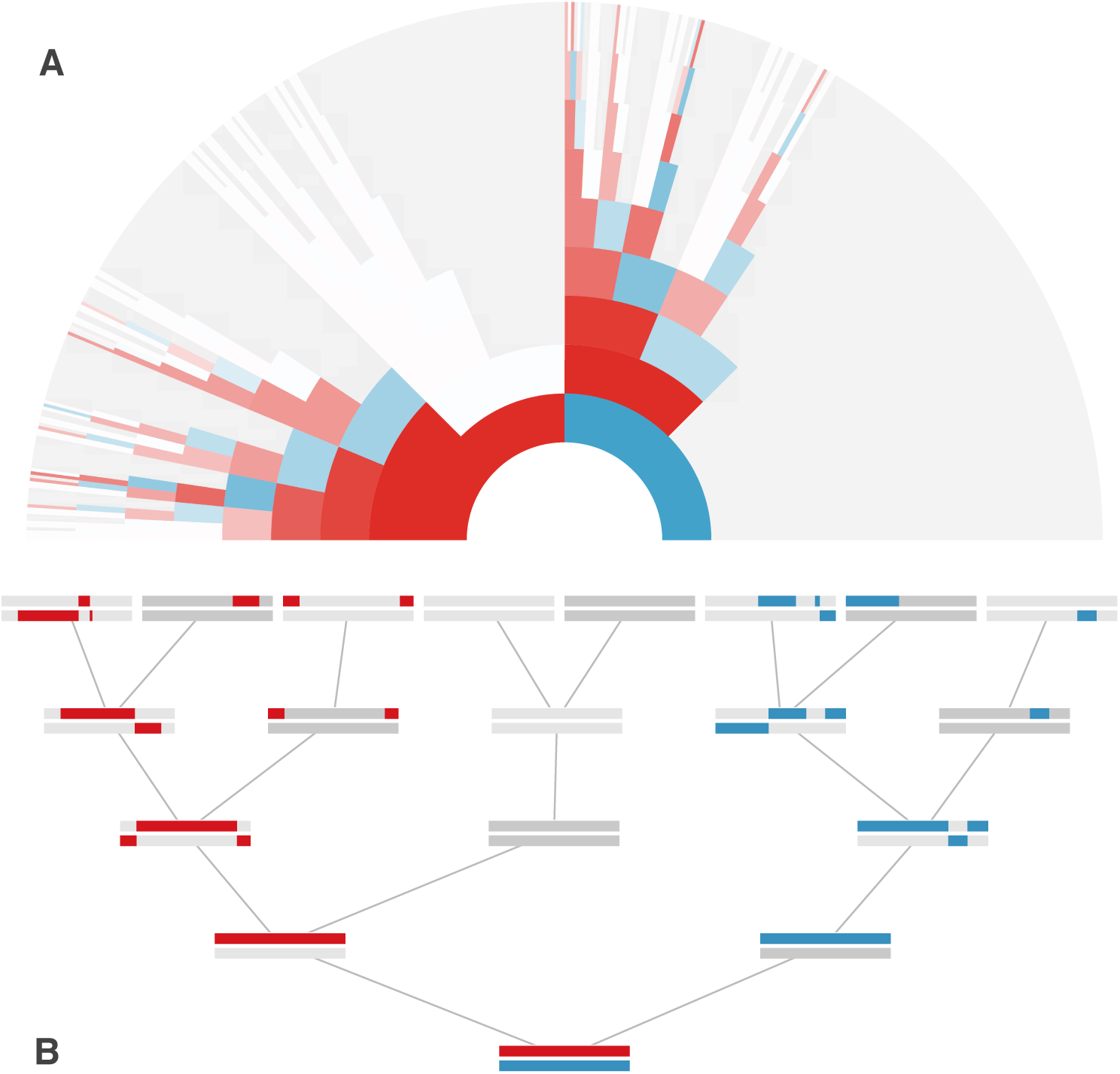
A: Simulated X genealogy of a present-day female, back nine generations. Each arc is an ancestor, with female ancestors colored red, and male ancestors colored blue. The transparency of each arc reflects the genetic contribution of this ancestor to the present-day female. White arcs correspond to an X genealogical ancestor that shares no genetic material with the present-day female, and gray arcs are genealogical ancestors that are not X ancestors. B: The X segments of the simulation in (A), back five generations. The maternal X lineage’s segments are colored red, and the paternal X segments are colored blue. A male ancestor’s sex chromosomes are colored dark gray (and include the Y) and a female ancestor’s sex chromosomes are colored light gray.

### The number of recombinational meioses along an unknown X lineage

If we pick an ancestor at random *k* generations ago, the probability that they are an X genealogical ancestor is given by equation (7). We can now extend this logic and ask: having randomly sampled an X genealogical ancestor, how many recombinational meioses (i.e. females) lie in the lineage between a present day individual and this ancestor? Since IBD segment number and length distributions are parameterized by a rate proportional to the number of recombination events, this quantity is essential to our further derivations.

Specifically, if there’s uncertainty about the particular lineage between a present-day female and one of her X ancestors *k* generations back (such that all of the *Ƒ*_*k*+2_ lineages to an X ancestor are equally probable), the number of females (thus, recombinational meioses) that occur is a random variable *R.* By the no two adjacent males condition, the possible number of females *R* is constrained; *R* has a lower bound of 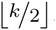, which corresponds to a male alternating each generation to an ancestor in the *k*^th^ generation. Similarly, the upper bound of *R* is *k*, since it is possible every individual along one X lineage is a female. Noting that an X genealogy extending back *k* generations enumerates every possible way to arrange *r* females such that none of the *k* – *r* males are adjacent, we find that the number of ways of arranging *r* such females this way is

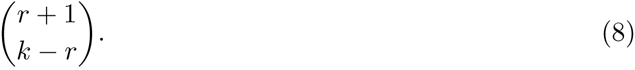

For some readers, it may be useful to visualize the relationship between the numbers of recombinational meioses across the generations using a variant of Pascals triangle (Figure 4). The sequence of recombinational meioses is related to a known integer sequence; see OEIS A030528 (Sloane, 2010) for a description of this sequence and its other applications.

**Figure 4:**
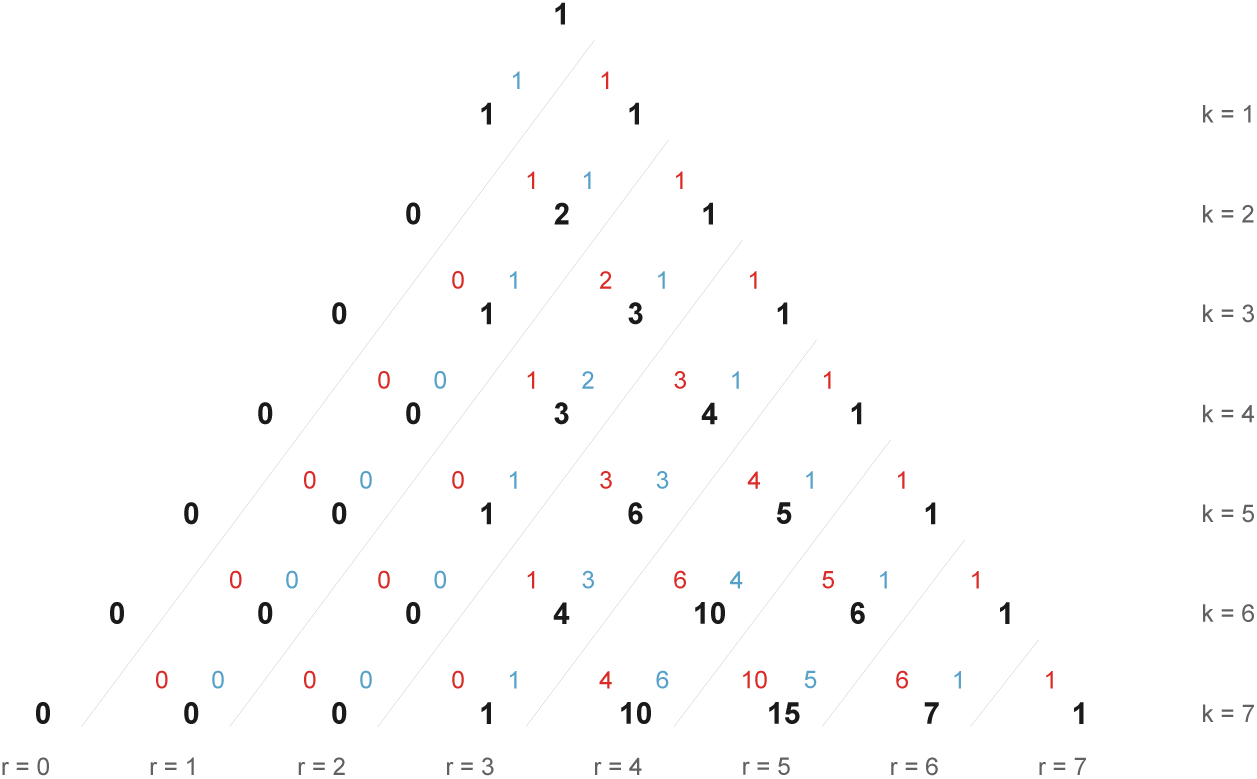
An triangle of 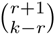 binomial coefficients, similar to an offset version of Pascal’s triangle. Black numbers indicate the number” of genealogical ancestors at generation *k* whose X chromosome would have been transmitted to the current generation by a route involving *r* recombinant meioses. Each value is further broken down into female (red value, upper left) and male (blue value, upper right) ancestors. Values for a new row in the triangle are generated by summing the black numbers to the left in the row above (these are the mothers of all ancestors in the next generation) and two rows directly above (these are male ancestors). The sum of each row (fixed *k*) is a Fibonacci number and the values in the diagonal corresponding to a fixed value of *r* are binomial coefficients.

If we pick an X genealogical ancestor at random *k* generations ago the probability that there are *r* female meioses along the lineage leading to this ancestor is

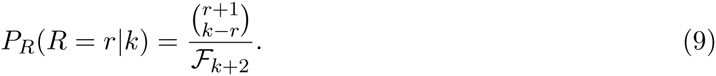

In Appendix B, we derive a generating function for the number of recombinational meioses. We can use this generating function to obtain properties of this distribution such as the expected number of recombinational meioses. We can show that the expected number of recombinational meioses converges rapidly to 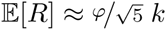 with increasing *k*, where *φ* is the Golden Ratio, 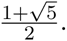

### The Distribution of Number of Segments Shared with an X Ancestor

Using the distribution of recombinational meioses derived in the last section, we now derive a distribution for the number of IBD segments shared between a present-day individual and an X ancestor in the *k*^th^ generation. For clarity, we first derive the number of IBD segments counted in the *parents* (i.e. not following the convention described in Section 1), but we can adjust this simply by replacing *k* with *k* – 1.

First, we calculate the probability of a present-day individual sharing *N* IBD segments with an X genealogical ancestor *k* generations in the past, where it is *known* that there are *R* = *r* females (and thus recombinational meioses) along the lineage to this ancestor. This probability uses the Poisson-Binomial model described in Section 1:

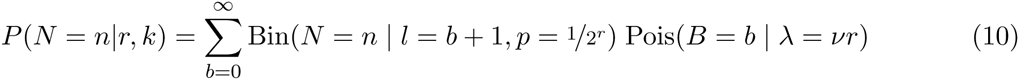

If we consider a single X genealogical ancestor *k* generations back, this individual could be any of the present-day female’s *Ƒ*_*k*+2_ X ancestors. Since the particular lineage to this ancestor is unknown, we marginalize over all possible numbers of recombinational meioses that could occur:

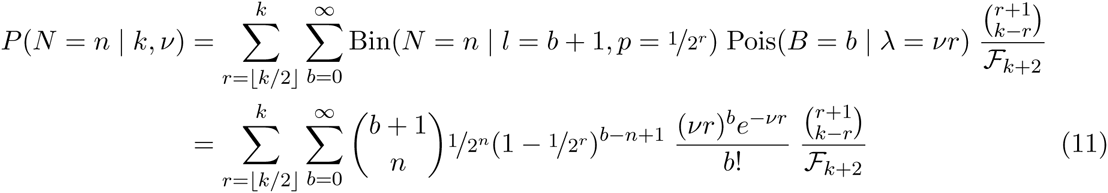

Note that once we’ve conditioned on the number of recombinational meioses *r*, the lineages to an X ancestor are interchangeable; the specific X lineage affects recombination (and thus the IBD number and length distributions) only through the number of recombinational meioses along the lineage.

For the distribution of number of IBD segments counted in the offspring, we substitute *k* – 1 for *k*

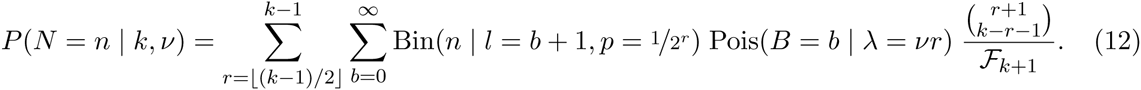

In this formulation, if *k* = 1, *r* = 0. In this case, the lack of recombinational meioses implies *b* = 0, such that a present-day female shares *n* = 1 X chromosomes with each of her two parents in the *k* = 1 generation with certainty.

We can use our equation (12) to obtain *P*(*N* > 0), the probability that a genealogical X ancestor *k* generations ago is a *genetic* ancestor. This probability over *k* ∈ {1, 2,…, 14} generations is shown in Figure 3B. For comparison, Figure 3B also includes the probability of a genealogical ancestor in the *k*^th^ generation being an autosomal genetic ancestor and the probability of being a genetic X ancestor unconditional on being an X genealogical ancestor.

We have also assessed the Poisson thinning approach to modeling X IBD segment number. As with the Poisson-Binomial model, we marginalize over *R:*

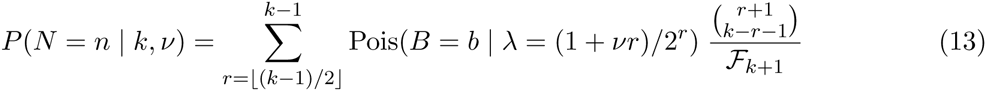

In Figure 5 we have compared the Poisson-Binomial and Poisson-thinning approximations for the number of IBD segments (counted in the offspring) shared between an X-ancestor in the *k*^th^ generation and a present-day female. Overall, the analytic approximations are close to the simulation results, with the Poisson-Binomial model a closer approximation for small *k* and both models’ accuracy improving quickly with increasing *k*. The Poisson-Binomial model is more accurate than the Poisson thinning approximation due to the overdispersion discussed in Section 1. As *k* increases, this extra variance quickly decreases. Note that since the total genetic length of the X is relatively short compared to the autosomes, the extra variance makes a relatively small contribution to the total variance (see Appendix equation (36)); this is why the Poisson thinning approach is a better approximation for autosomal IBD segments. See Appendix A for details. Throughout the paper, we use the more accurate Poisson-Binomial model rather than this Poisson thinning model. If only X ancestry more than 3 generations back is of interest, the Poisson thinning approach may be used without much loss of accuracy.

**Figure 5:**
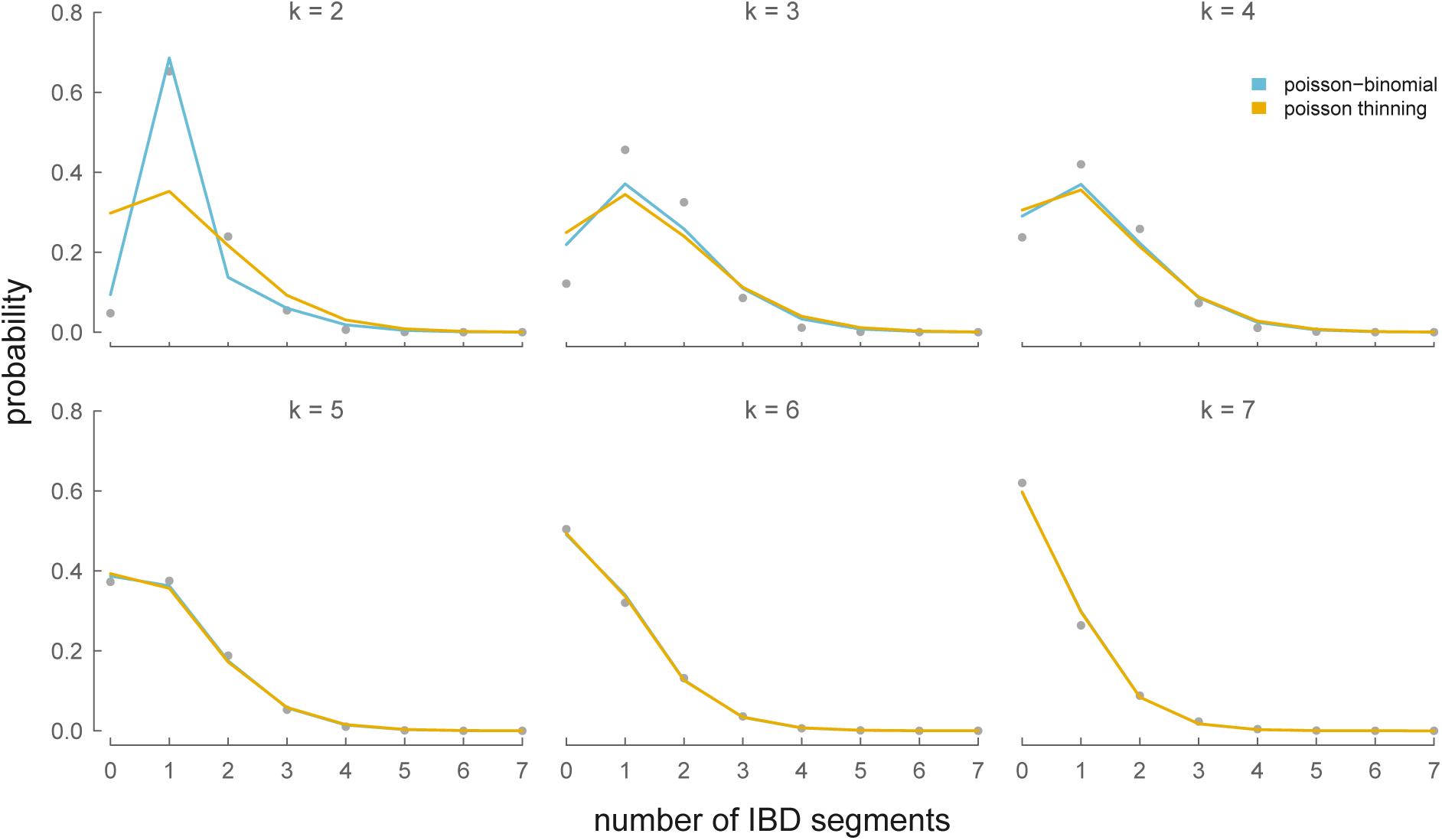
The Poisson thinning (yellow lines) and Poisson-Binomial (blue lines) analytic distributions of IBD segment number between an X ancestor in the *k*^th^ generation (each panel) and a present-day female. Simulation results averaged over 5,000 simulations are the gray points.

### The Distribution of IBD Segment Lengths with an X Ancestor

The distribution of IBD segment lengths between a present-day female and an unknown X genealogical ancestor in the *k*^th^ generation is similar to the autosomal length distribution (equation 6). However, with uncertainty about the particular lineage to the X ancestor, the number of recombinational meioses can vary between 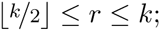 we marginalize over the unknown number of recombinational meioses using the distribution equation (9). Our length density function is:

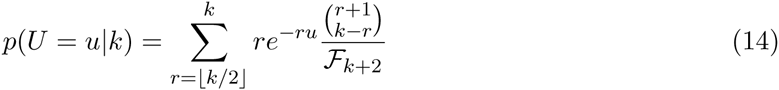

In Figure 6, we compare our analytic length density to an empirical density of X segment lengths calculated from 5,000 simulations. As with our IBD segment number distributions, our analytic model is close to the simulated data’s empirical density, and converges rapidly with increasing *k*.

**Figure 6:**
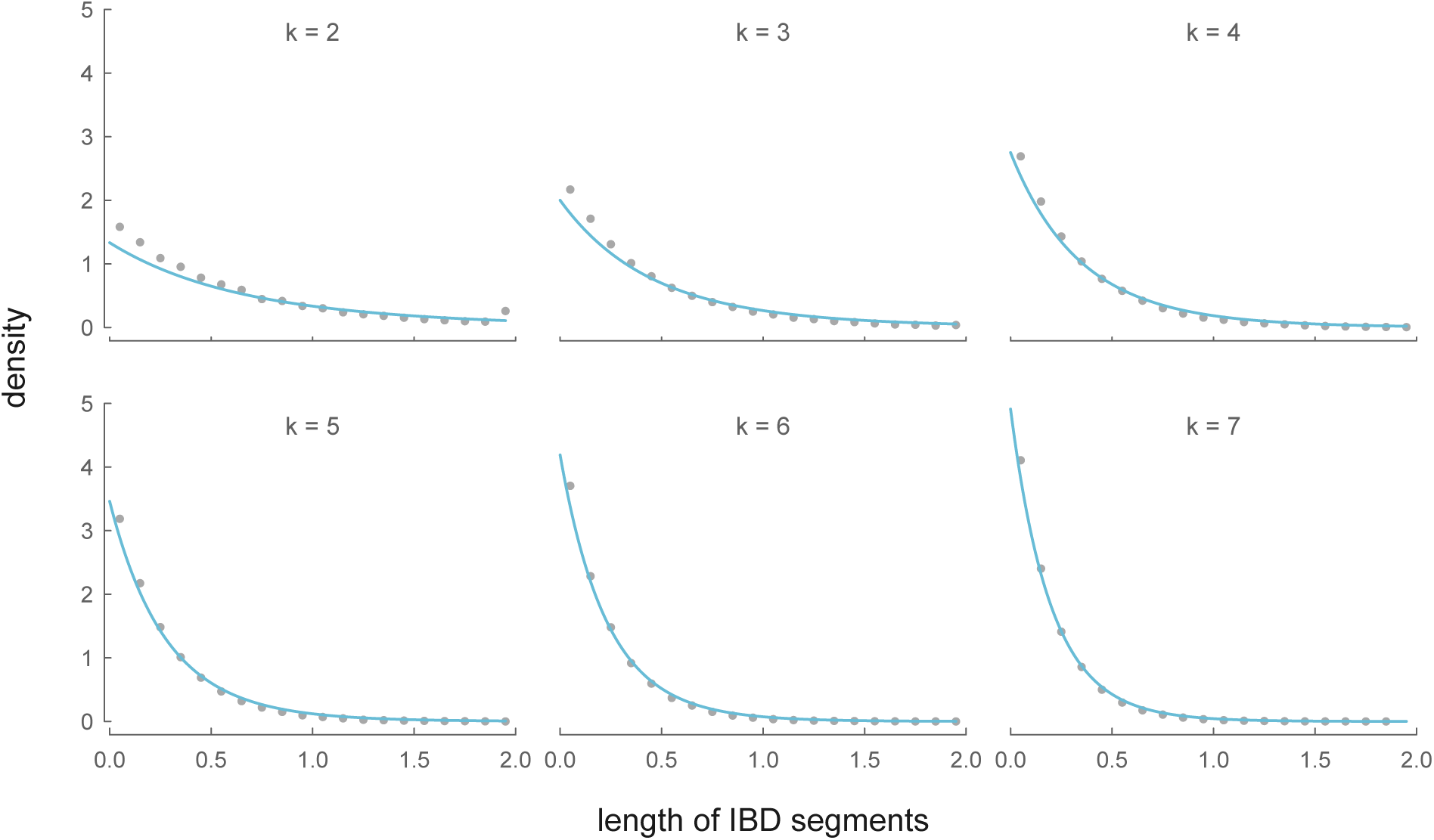
The analytic distributions of IBD segment length between an ancestor in the *k*^th^ generation (for *k* ∈ {2,…, 7}) and a present-day female (blue lines), and the binned average over 5,000 simulations (gray points).

Note that both the IBD segment length and number distributions marginalize over an unobserved number of recombinational meioses (*R*) that occur along the lineage between individuals. As the IBD segments shared between two individuals is a function of the number breakpoints *B*, and thus recombinational meioses, neither the length nor number distributions *P*(*N* = *n*) and *p*(*U* = *u*) (which separately marginalize over both *R* and *B*) are independent of each other. Intuitively, we can understand this dependence in the following way: if we observe one long segment nearly the entire length of the X, this makes observing a large number of additional segments in the remaining region of the X unlikely.

## 3 Shared X Ancestry

Because only a fraction of one’s genealogical ancestors are X ancestors (and this fraction rapidly decreases with *k*; see equation (7)), two individuals sharing X segments IBD from a recent ancestor considerably narrows the possible ancestors they could share. In this section, we describe the probability that a genealogical ancestor is an X ancestor, and the distributions for IBD segment number and length across full-and half-cousin relationships. For simplicity we concentrate on the case where the cousins share a genealogical ancestor *k* generations ago in both of their pedigrees, i.e. the individuals are *k* – 1 degree cousins. The formulae could be generalized to ancestors of unequal generational-depths (e.g. second cousins once removed) but we do not pursue this here.

### Probability of a Shared X Ancestor

Given that two individuals share their first common genealogical ancestor in the *k*^th^ generation, we can calculate the probability that this single ancestor is also an X genealogical ancestor. We can think of this first shared ancestor in the *k*^th^ generation as the same individual in both of the two sets of 2^k^ ancestors of each present-day individual. Since this shared ancestor must be of the same sex in each of the two present-day individuals’ genealogies, we condition on the ancestor’s sex (with probability 1/2 each) and then calculate the probability that this individual is also an X ancestor (with the same sex). Let us define 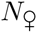 and 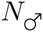 as the number of genealogical female and male ancestors, and 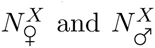 as the number of X female and male ancestors of a present-day individual in the *k*^th^ generation. Then:

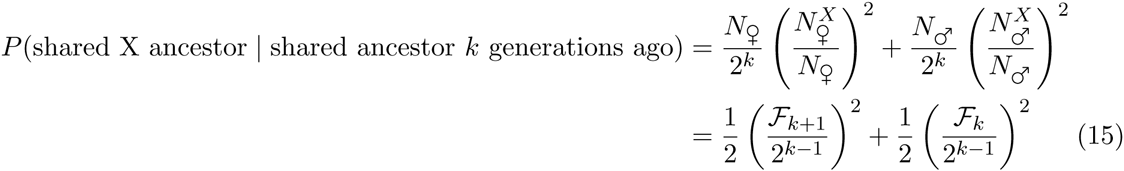

Thus, the probability that a shared genealogical ancestor is also a shared X ancestor is decreasing at an exponential rate. By the 8^th^ generation, a shared genealogical ancestor has less than a five percent chance of being a shared X ancestor of both present-day individuals.

### The Sex of Shared Ancestor

Unlike genealogical ancestors, which have equal numbers of female and male ancestors, recent X genealogical ancestors are predominantly female. Since a present-day female has *Ƒ*_*k*+1_ female ancestors and *Ƒ*_*k*_ male ancestors *k* generations ago, the ratio of female to male X genealogical ancestors converges to the Golden Ratio 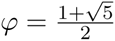 (Simson, 1753; Wells, 1997):

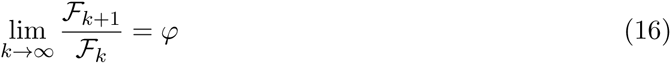

In modeling the IBD segment number and length distributions between present day individuals, the sex of the shared ancestor *k* generations ago affects the genetic ancestry process in two ways. First, a female shared ancestor allows the two present-day individuals to share segments on either of her two X chromosomes while descendents of a male shared ancestor share IBD segments only through his single X chromosome. Second, the no two adjacent males condition implies a male shared X genealogical ancestor constrains the X genealogy such that the present-day X descendents are related through his two daughters. Given that the ratio of female to male X ancestors is skewed, our later distributions require an expression for the probability that a shared X ancestor in the *k*^th^ generation is female, which we work through in this section.

As in equation (15), an ancestor shared in the *k*^th^ generation of two present-day individuals’ genealogies must have the same sex in each genealogy. Assuming both present-day cousins are females, in each genealogy there are *Ƒ*_k_ possible male ancestors and *Ƒ*_*k*+1_ female ancestors that could be shared. Across each present-day females’ genealogies there are (*Ƒ*_*k*_)^2^ possible male ancestor combinations and (*Ƒ*_*k*+1_)^2^ possible female ancestor combinations. Thus, if we let 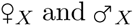 denote that the sex of the shared is female and male respectively, the probability of a female shared ancestor is:

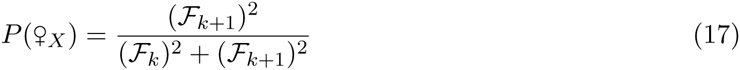

The probability that the shared ancestor is male is simply 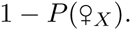 One curiosity is that as 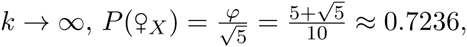, where *φ* is the Golden Ratio.

### Partnered Shared Ancestors

Thus far, we have only looked at two present-day individuals sharing a single X ancestor *k* generations back. In monogamous populations, most shared ancestry is likely to descend from *two* ancestors; we call such relationships *partnered* shared ancestors. In this section, we look at full-cousins descending from two shared genealogical ancestors that may also be X ancestors. Two full-cousins could either (1) both descend from two X ancestors such that they are X full-cousins, (2) share only one X ancestor, such that they are X half-cousins, or (3) share no X ancestry. We calculate the probabilities associated with each of these events here.

Two individuals are full-cousins if the great^*k*–2^ grandfather and the great^*k*–2^ grandmother in one individual’s genealogy are the same couple as in the other individual’s genealogy. For these two full-cousins to be X full-cousins, this couple must also be a couple in both individuals’ X genealogies. In every X genealogy, the number of couples in generation *k* is the number of females in generation *k* – 1, as every female has two X ancestors in the next generation (while males only have one). Thus, the probability two female *k* – 1 degree full-cousins are also X full-cousins is:

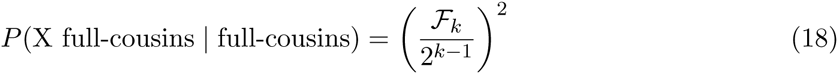

Now, we consider the event that two genealogical full-cousins are X half-cousins. Being X half-cousins implies that the partnered couple these full-cousins descend from includes a single ancestor that is in the X genealogies of both full-cousins. This single X ancestor must be a female, as a male X ancestor’s female partner must also be an X ancestor (since mothers must pass an X). For a female to be an X ancestor but not her partner, one or both of her offspring must be male. Either of these events occurs with probability:

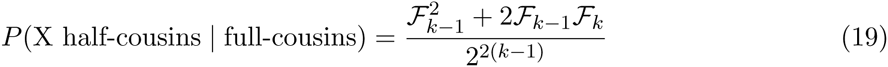

### The Distribution of Recombinational Meioses between Two X Half-Cousins

To find distributions for the number and lengths of IBD segments shared between two half-cousins on the X chromosome, we first need to find the distribution for the number of females between two half-cousins with a shared ancestor in the *k*^th^ generation. We refer to the individuals connecting the two cousins as a *genealogical chain*. As we’ll see in the next section, the number of IBD X segments shared between half-cousins depends on the sex of the shared ancestor; thus, we also derive distributions in this section for the number of recombinational meioses along a genealogical chain, conditioning on the sex of the shared ancestor. As earlier, our models assume two present-day female cousins but are easily extended to male cousins.

First, there are 2*k* – 1 ancestral individuals separating two present-day female (*k* – 1)^th^ degree cousins. These X ancestors in the genealogical chain connecting the two present-day female cousins follow the no two adjacent male condition; thus the distribution of females follows the approach used in equation (9) with *k* replaced with 2*k* – 1:

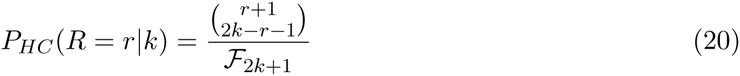
where the *HC* (for half-cousin) subscript differentiates this equation from equation (9) and *k* is the generation of the shared ancestor. Similarly to equation (9), *R* is bounded such that ⌊(2*k* – 1)/2⌋ ≤ *R* ≤ 2*k* – 1.

Now, we derive the probability of *R* = *r* females conditional on the shared ancestor being female, ♀x. This conditional distribution differs from equation (20) since it eliminates all genealogical chains with a male shared ancestor.

We find the distribution of recombinational meioses conditional on a female shared ancestor by placing the other *R′* = *r′* females (the prime denotes we do not count the shared female ancestor here) along the two lineages of *k* – 1 individuals from the shared female ancestor down to the present-day female cousins. These *R′* = *r′* females can be placed in both lineages by positioning *s* females in the first lineage and *r′* – *s* females in the second lineage, where *s* follows the constraint ⌊(*k* – 1)*/*2⌋ ≤ *s* ≤ (*k* - 1)/2. Our equation (9) models the probability of an X genealogical chain having *r* females in *k* generations; here, we use this distribution to find the probabilities of *s* females in *k* – 1 generations in one lineage and *r′* – *s* females in *k* – 1 generations in the other lineage. As the number of females in each lineage is independent, we take the product of these probabilities and sum over all possible *s*; this is the discrete convolution of the number of females in two lineages *k* – 1 generations long. Finally, we account for the shared female ancestor, by the transform *R* = *R′*+ 1 = r:

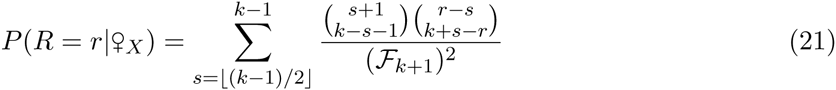

In general, this convolution approach allows us find the distribution of females in a genealogical chain under various constraints, and can easily be extended to the case of a shared male X ancestor (with necessarily two daughters).

Finally, note that we have modeled the number of *females* in a genealogical chain of *2k*– 1 individuals. Thus far in our models, the number of females has equaled the number of recombinational meioses. However, when considering the number of recombinational meioses between half-cousins, *two* recombinational meioses occur if the shared ancestor is a female (as she produced two independent gametes she transmits to her two offspring). Thus, for a single shared X ancestor, the number of recombinational meioses *ρ* is

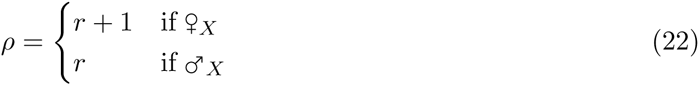
which we use when parameterizing the rate of recombination in our IBD segment number distributions. Furthermore, since a shared female ancestor has two X haplotypes that present-day cousins could share segments IBD through, the binomial probability 1/2^*ρ*^ is doubled. Further constraints are needed to handle full-cousins; we will discuss these in Section 3.

### Half-Cousins

In this section we calculate the distribution of IBD X segments shared between two present-day female X half-cousins with a shared ancestor in the *k*^th^ generation. We imagine we do not know any details about the lineages to this shared ancestor nor the sex of the shared ancestor, so we marginalize over both. Thus, the probability of two (*k* – 1)^th^ degree X half-cousins sharing *N* = *n* segments is:

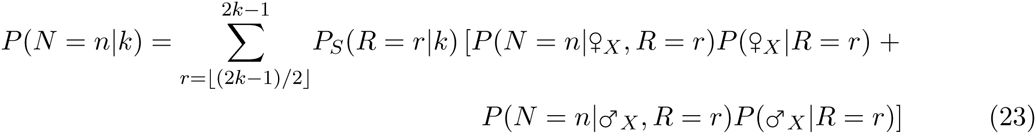

As discussed in the previous section, the total number of recombinational meioses along the genealogical chain between half-cousins depends on the unobserved sex of the shared ancestor (i.e. equation (22)). Likewise, the binomial probability also depends on the shared ancestor′s sex. Accounting for these adjustments, the probabilities 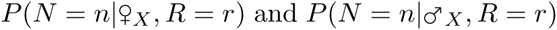 are:

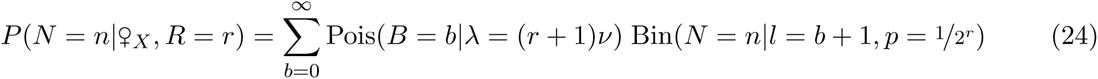

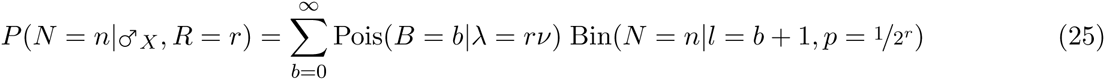

Since the sex of the shared ancestor depends on the number of females in the genealogical chain between the two cousins (e.g. if *r* = 2*k* – 1, the shared ancestor is a female with certainty), we require an expression for the probability of the shared ancestor being male or female given *R* = *r*. Using Bayes’ theorem, we can invert the conditional probability *P*(*R* = *r*|♀*X*) to find that the probability that a shared X ancestor is female conditioned on *R* females in the genealogical chain is

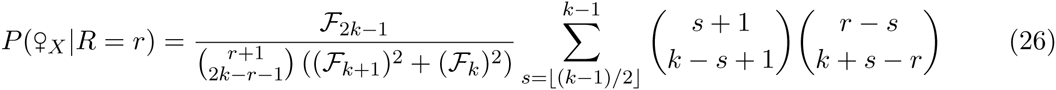
and *P*(♂_*X*_|*R* = *r*) can be found as the complement of this probability.

Inserting equations (25), (24), and (26) into (23) gives us an expression for the distribution of IBD segment numbers between two half-cousins with a shared ancestor *k* generations ago. Figure 7 compares the analytic model in equation (23) with the IBD segments shared between half-cousins over 5,000 simulated pairs of X genealogies.

**Figure 7:**
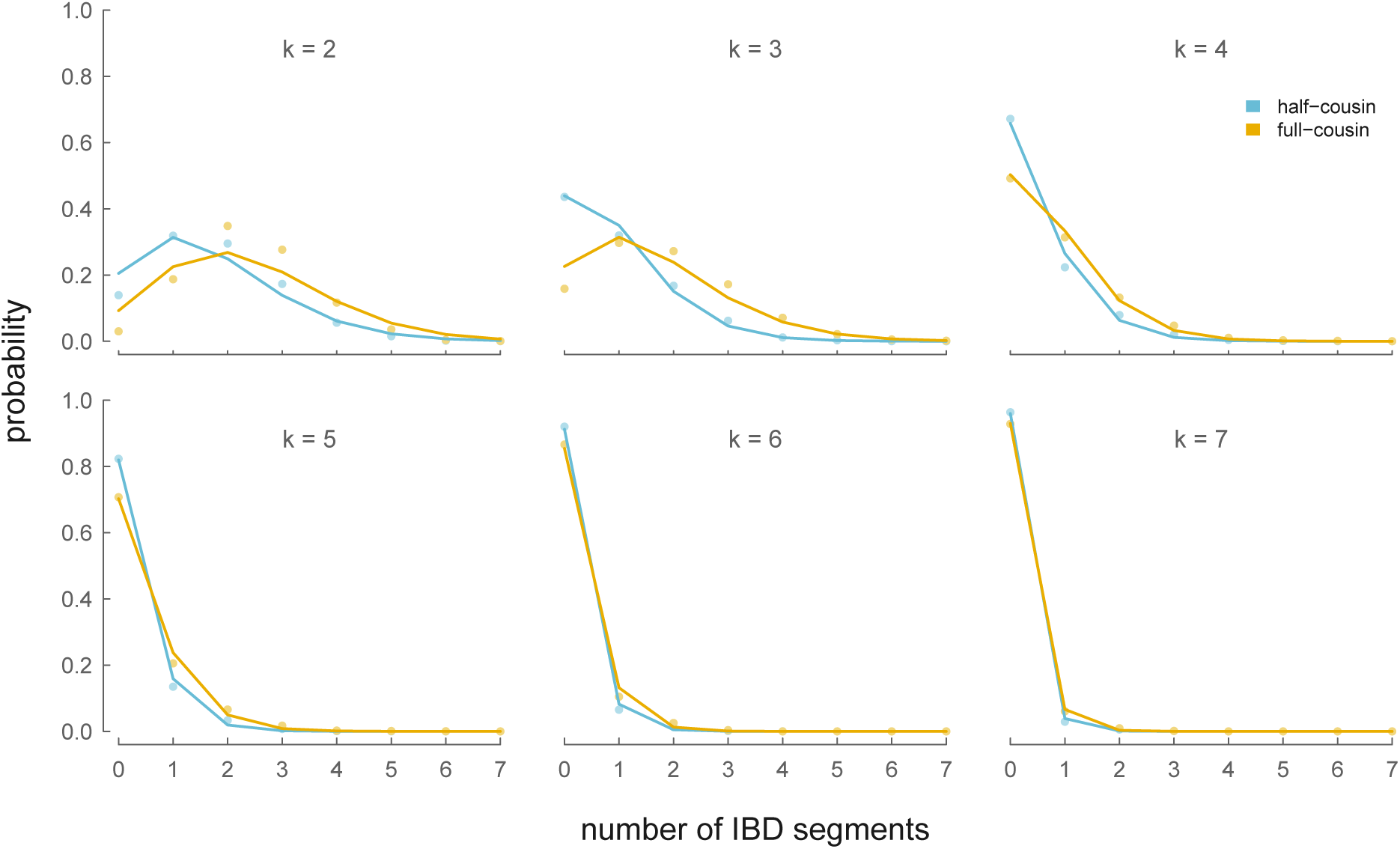
Distributions of X IBD segment number for X half-(blue) and X full-cousins (yellow). Lines show the analytic approximations (equations (23) and (28)) and gray points show the probability averaged over 5,000 simulations.

The density function for IBD segment lengths between X cousins (either half-or full-cousins; length distributions are only affected by the number of recombinations in the genealogical chain) is equation (14) but marginalized over the number of recombinational meioses between two cousins (equation (20)) rather than the number of recombinational meioses between a present-day individual and a shared ancestor. Simulations show the length density closely matches simulation results (see Figure 11 in the appendix).

**Figure 11:**
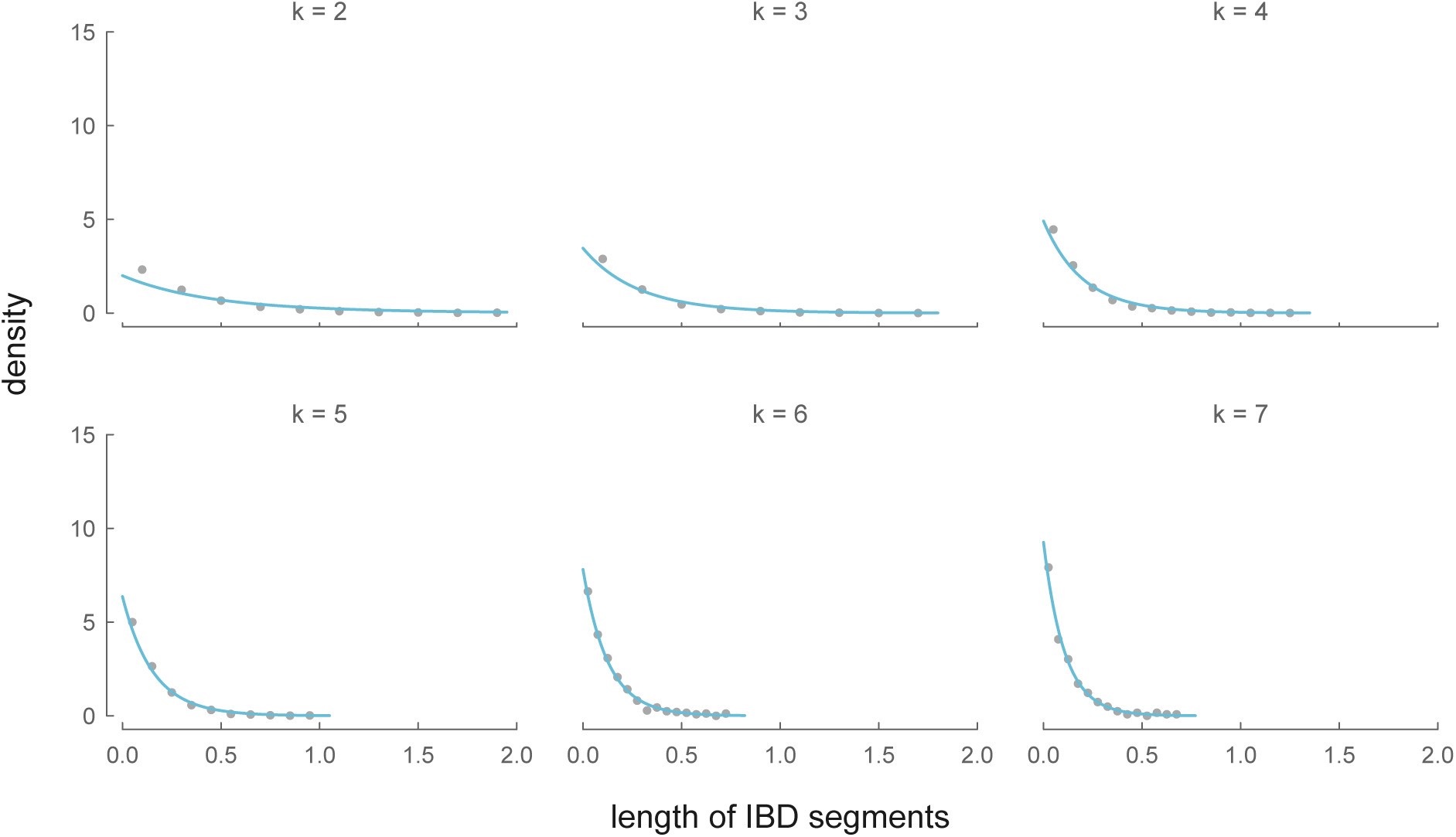
The analytic distributions of IBD segment length (blue lines) between two-present-day female half-cousins with a shared ancestor in the *k*^th^ generation (each panel), and the binned average 5,000 simulations (gray points).

### Full-Cousins

Full-cousin relationships allow descendents to share IBD autosomal segments from either their shared maternal ancestor, shared paternal ancestral, or both. In contrast, since males only pass an X chromosome to daughters, only full-sibling relationships in which both offspring are female (due to the no to adjacent males condition) are capable of leaving X genealogical descendents. We derive a distribution for the number of IBD segments shared between (*k* – 1)^th^ degree full X cousins by conditioning on this familial relationship and marginalizing over the unobserved number of females from the two full-sibling daughters to the present-day female full-cousins.

First, we find the number of females (including the two full-sibling daughters in the (*k* – 1)^th^ generation) in the genealogical chain between the two full-cousins (omitting the shared male and female ancestors, which we account for separately). Like equation (21), this is a discrete convolution:

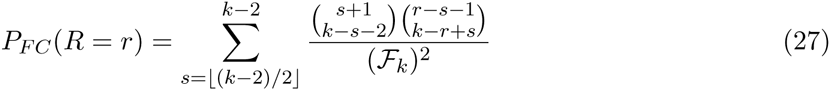

This probability is valid for 2⌊(*k* - 2)/2⌋ + 2 ≤ *r* ≤ 2*k* – 2 and is 0 elsewhere. For *N* = *n* segments to be shared between two full-cousins, *z* segments can be shared via the maternal shared X ancestor (where 0 ≤ *x* ≤ *n*) and *n* – *z* segments can be shared through the paternal shared X ancestor. We marginalize over all possible values of *z*, giving us another discrete convolution:

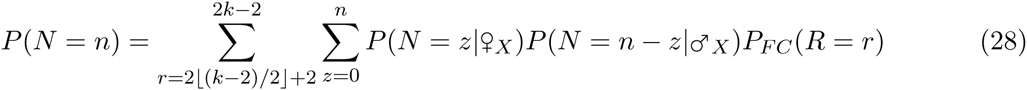

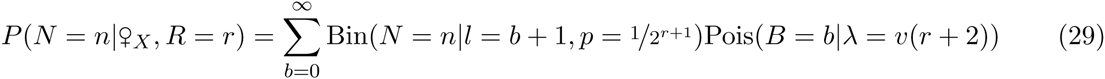

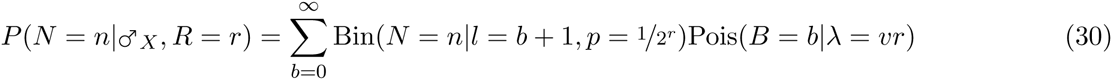
are the probabilities of sharing *n* segments through the shared female and male X ancestors respectively. For the female shared ancestor, we account for two additional recombinational meioses (one for each of the two gametes she passes to her two daughters), and the fact she can share segments through either of her X chromosomes (hence, why the binomial probability is 1/2^*r*+1^). We compare our analytic X full-cousin IBD segment number results to 5,000 genealogical simulations in Figure 7.

## 4 Inference

With our IBD X segment distributions, we now turn to how these can be used to infer recent X ancestry. In practice, inferring the number of generations back to a common ancestor (*k*) is best accomplished through the signature of recent ancestry from the 22 autosomes, rather than through the short X chromosome. Therefore, we concentrate on questions about the extra information that the X provides conditional on *k* being known with certainty. A number of methods are available for the task of estimating *k* through autosomal IBD segments (Durand et al., 2014; Henn et al., 2012; Huff et al., 2011).

Here, we focus on two separate questions: (1) what is the probability of being an X genealogical ancestor given that no IBD segments are observed, and (2) can we infer details about the X genealogical chain between two half-cousins? These questions address how informative the number of segments shared between cousins is about the precise relationship of cousins. We assume that segments of X chromosome IBD come only from the *k*^th^ generation, and not from deeper relationships or from false positives. In practice, inference from the X IBD segments would have to incorporate both of these complications, and as such our results represent best case scenarios.

It’s possible that *k* generations back, an individual is a genealogical X ancestor but shares no X genetic material with a present-day descendent. To what extent is the lack of sharing on the X chromosome with an ancestor informative about our relationship to them? How does the lack of sharing of the X chromosome between *k* – 1 cousins change our views as to their relationship? To get at these issues, we can use our analytic approximations to calculate the probability that one is an X ancestor given that no segments are observed, *P*(X ancestor | *N* = 0):

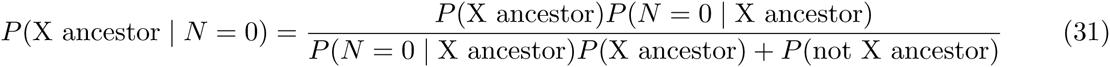

Here, *P*(*N* = 0 | X ancestor) is given by equation (12) and *P*(X ancestor) is given by equation (7). This function is shown in Figure 8 (yellow lines). We can derive an analogous expression for the probability of two female half-cousins sharing an X ancestor but not having any X segments IBD by replacing *P*(X ancestor |*N* = 0) with equation (23), and replacing *P*(X ancestor) with *P*(shared X ancestor) which is given by equation (15) (blue lines). We also plot the prior distributions to show the answer if no information about the X chromosome was observed. In both cases, the lack of sharing on the X chromosome does narrow the field of ancestors, making those who are not X genealogical ancestors more likely, beyond our prior. This is especially true for close relationships (*k* < 5), where the X is likely to be shared if the ancestor was an X genealogical ancestor.

**Figure 8:**
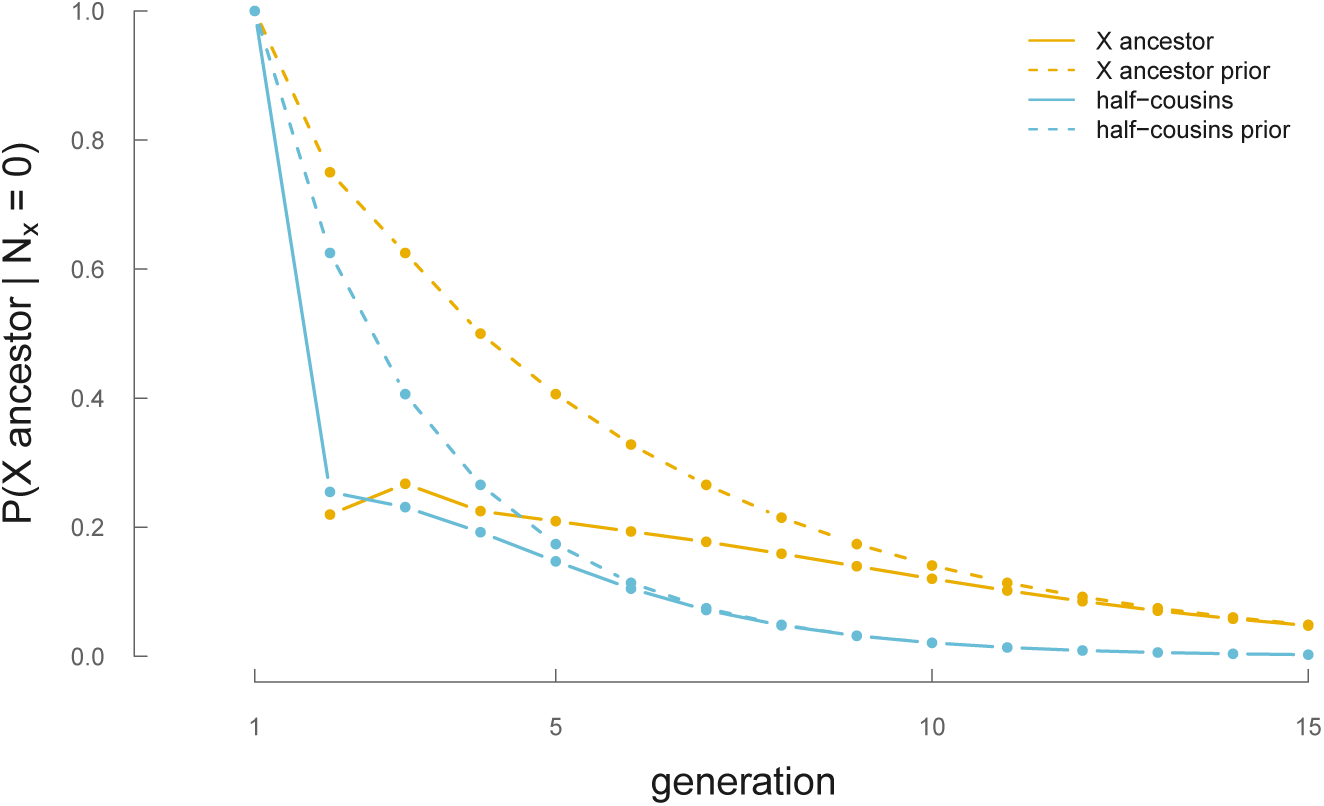
The probabilities of an individual being an X ancestor to a present-day female (yellow solid line) or single shared X ancestor (blue solid line) to two present-day female cousins conditional on not sharing any X IBD segments, *k* generations back. Dashed lines indicate the prior probabilities.

X IBD segments can be used to infer genealogical details that are not possible looking only at autosome IBD segments. Unlike the X, the autosomes undergo recombinational meioses in every individual each generation. Consequently, while IBD autosome segments leave a signature of recent ancestry between two individuals, the uniformity of recombinational meioses across every lineage to the shared ancestor leaves no signal of *which* genealogical lineage connects two present-day cousins.

In contrast, X genealogical chains have two properties that leave a signature of which genealogical lineage connects two present-day cousins. First, recombinational meioses only occur in females. Consequently, more females along an X genealogical chain increase the rate of recombination and alter the IBD segments length and number distributions in a lineage-specific manner. Second, the number of females varies across X lineages; for example the number of females between two present-day half-cousins varies by a factor of two or more. These two properties allow us to find a posterior distribution for the number of females that separate two present-day cousins. This number of females *R* is our only inferable summary of the genealogical chain and constrains the possible X genealogical chains between these two cousins by varying amounts dependent on *R* and *k*.

Our approach to inference is through the posterior distribution of *R* given an observed number of IBD segments *N* and conditioning on *k*. We calculate this posterior conditional on the cousins sharing an X ancestor; we do this to separate it from the question of whether a pair share an X ancestor (as derived in equation (31)). Our posterior probability is given by Bayes’ theorem

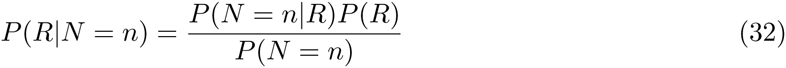
where the prior *P*(*R*) is readily calculable through equation (20) and *P*(*N* = *n*) given by equation (23). The data likelihood *P*(*N* = *n*\*R*) is given by equation (10).

In Figure 9, we show the posterior distributions over the number of recombinational meioses, given an observed number of IBD segments between two females known to be X half-cousins. Again, these posterior distributions condition on knowing how many generations have occurred since the shared ancestor, *k.* With an increasing number of generations to the shared ancestor, fewer segments survive to be IBD between the present-day cousins. Consequently, observing IBD segments increases the likelihood of fewer females (and thus fewer recombinational meioses) between the cousins. For example, for *k* = 6, observing (the admittedly unlikely) six or more IBD segments leads to a posterior mode over the fewest possible number of females in the genealogical chain (⌊(2*k* – 1)/2⌋ = 5; Figure 9). Similarly, observing between three and five segments places the posterior mode over six females in the genealogical chain. For *k* > 4, seeing zero segments provides little information over the prior about the relationship between the cousins, as sharing zero segments is the norm.

**Figure 9:**
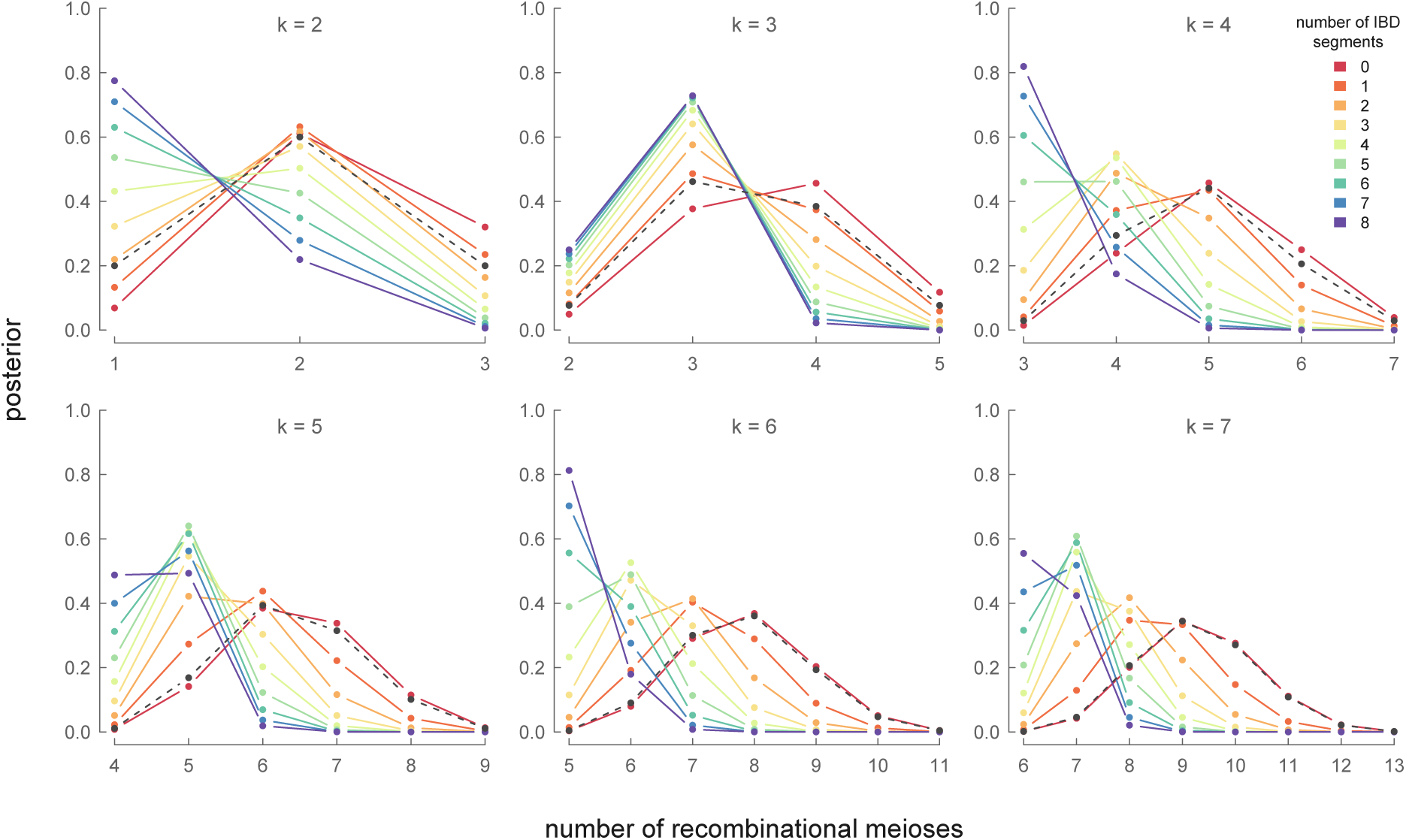
Posterior probability distribution *P*(*R* = *r*|*N* = *n, k*) for different generations (each panel), and the observed number of IBD segments (each colored line). The prior distribution of recombinational meioses given *k* is indicated by a gray dashed line.

## 5 Discussion

Detecting and inferring the nature of recent ancestry is important to a range of applications and the nature of such relationships are often of inherent interest. As the sample sizes of population genomic data sets increase, so will the probability of sampling individuals that share recent ancestry. In particular, the very large data sets being developed in human genetics will necessitate taking a genealogical view of recent relatedness. Our methods extend existing methods for the autosomes by accounting for the special inheritance pattern of the X. Specifically, recent ancestry on the X differs from the autosomes since males only inherit an X from their mothers, and fathers pass an unrecombined (ignoring the PAR) X to their daughters. Consequently, the number of recombinational meioses, which determine the length and number of IBD segments, varies across the X genealogy. Since in most cases the number of females between two individuals in a genealogical chain is often unknown, we derive a distribution for recombinational meioses (equation (9)).

We derive distributions for the length and number of IBD X segments by marginalizing over the unknown number of recombinational meioses that can occur between two individuals connected through a genealogical chain. In both cases, we condition on knowing *k* (the generations back to a shared ancestor) which can be inferred from the autosomes (Huff et al., 2011). Our models for IBD segment number and length use a Poisson-Binomial approximation to the recombination process, which match simulation results closely.

Our results here not only allow X IBD segments to be used to model recent ancestry, but are in fact more informative about which genealogical ancestors individuals share than autosomal data alone. This additional information occurs through two avenues. First, sharing IBD segments on the X immediately reduces the potential genealogical ancestors two individuals share, since one’s X ancestors are only a fraction of their possible genealogical ancestors (i.e. *Ƒ*_*k*+2_/2^*k*^ in the case of a present-day female). Second, the varying number of females in an X genealogy across lineages combined with the fact that recombinational meioses only occur in females to some extent leave a lineage-specific signature of ancestry.

Unfortunately, the X chromosome is short, such that the chance of any signal of recent ancestry on the X decays rather quickly. However, growing sample sizes will increase both the detection of the pairwise relatedness and cases of relatedness between multiple individuals. In these large data sets, overlapping pairwise relationships (e.g. a present-day individual that shares X segments with two distinct other individuals) could be quite informative about the particular ancestors that individuals share.

Our results should also be of use in understanding patterns of admixture on the X chromosome. In particular our results about the posterior information from the number and length of X segments shared with a genealogical ancestor can help us understand what can be learned from the presence (or absence) of particular segments of particular ancestry on the X chromosome. While this information for an individual decays somewhat quickly after a small number of generations, models of X chromosome segment ancestry will be useful at a population-level for understanding sex-biased admixture (Bryc et al., 2010; Goldberg et al., 2015; Shringarpure et al., 2016).

## 6 Acknowledgements

We wish to thank Nancy Chen, Jeremy Berg, Kristin Lee, and the rest of the Coop lab for helpful discussions and feedback on earlier drafts. This research was supported by an NSF Graduate Research Fellowship grant awarded to VB (1148897), and National Institute of General Medical Sciences of the National Institutes of Health under award numbers NIH RO1GM83098 and RO1GM107374 to GC.

## A Convergence of the Thinned Poisson Process to Poisson-Binomial Model

We compared the Poisson thinning approximation and the Poisson-Binomial models. One can show using the law of total expectation that the Poisson-Binomial and Poisson model have the same expected value:

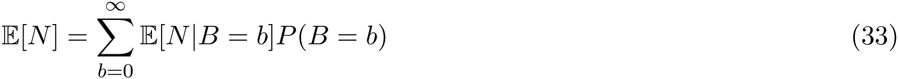

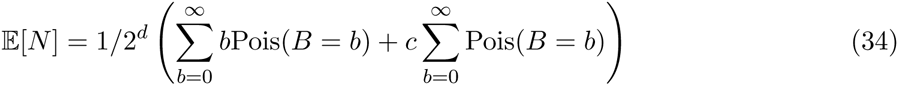

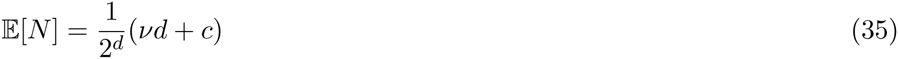

This is the same expected value as the thinned Poisson process with rate ( *vd* + *c*)/2^d^. However, the Poisson thinning and Poisson-Binomial models differ in their variance. Using Eve’s law, we can show the Poisson-Binomial model has variance

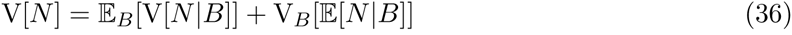

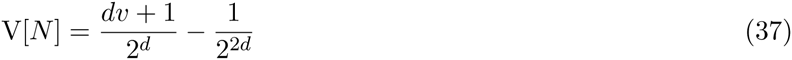

This differs from the thinned Poisson process variance by the term 1/2^2*d*^, which grows smaller with increasing *d.* Finally, we numerically show these two distributions (here, we label the two distributions for *k* generations *μ*_*k*_(*x*) and*v*_*k*_(*x*), where *x* is the number of segments) converge quickly in total variational distance 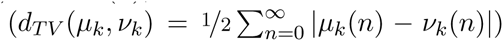 increases, in Figure 10.

**Figure 10:**
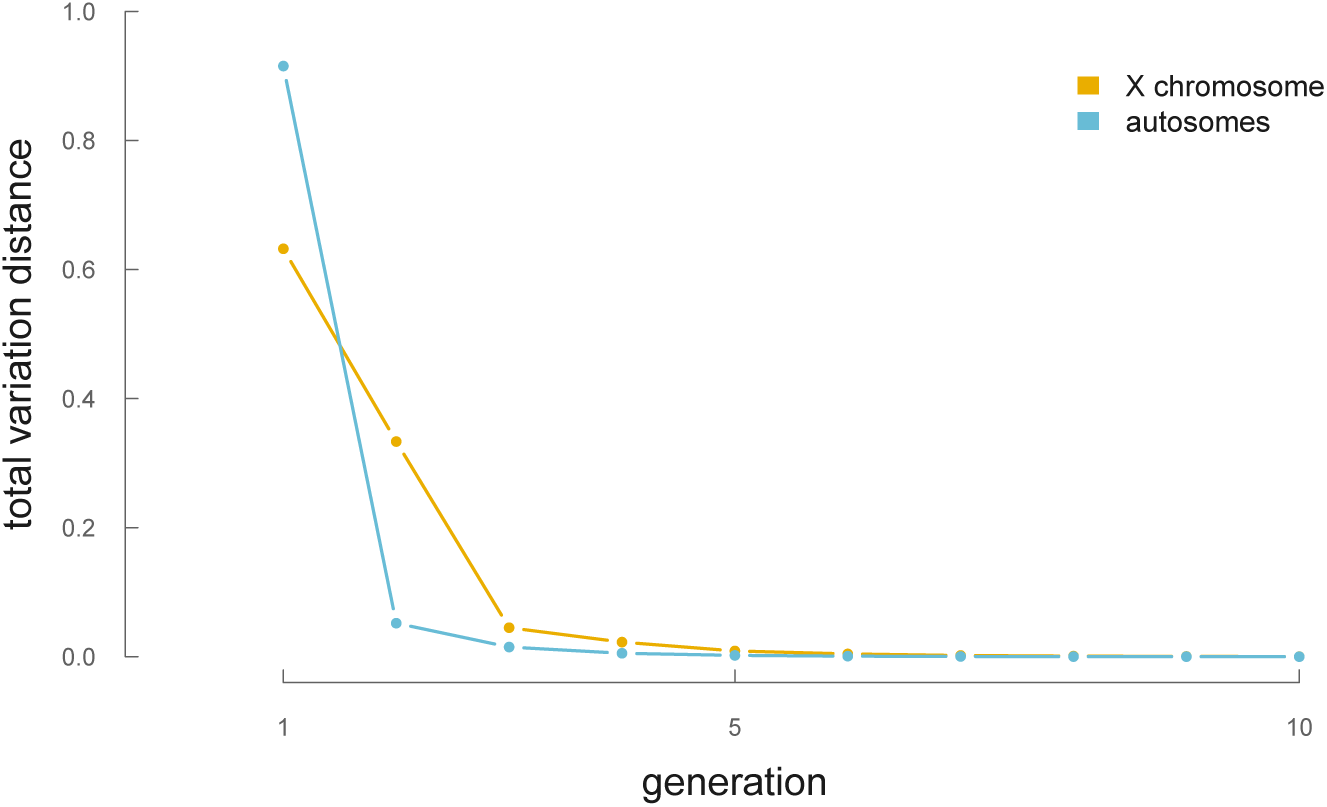
The total variation distance between the Poisson thinning and the Poisson-Binomial model for IBD segment number for X segments (yellow) and the autosomal segments (blue).

## B Generating Function for Recombinational Meioses

We also develop a generating function *g*(*x*, *k*) that encodes the number of recombinational meioses in the *k*^th^ generation as the coefficient for the term *x^k^.* This generating function can also be used in approximations and finding moments of the distribution *p*_k_(*r*).

**Lemma 1.** *An expansion of the generating function below encodes the number of lineages with r females in a genealogical chain k generations long (n_k,r_) as the coefficient of the term x^k^*

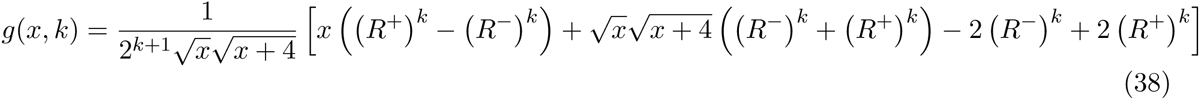

where

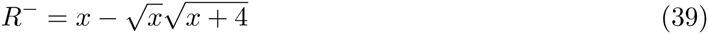

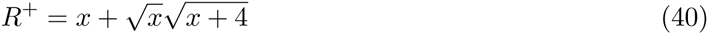

*Proof.* We begin by stating some recurrences that occur from the inheritance pattern of X ancestry:

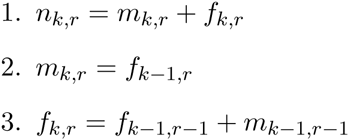

Starting from (1):

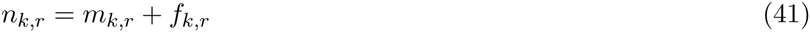

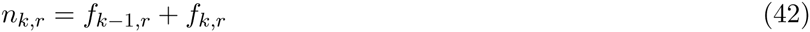

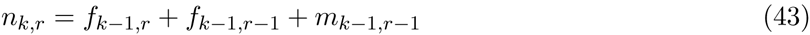

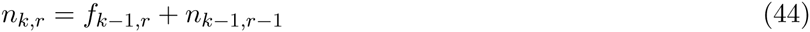

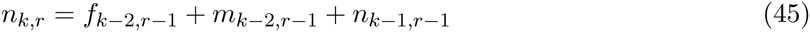
finally, substituting (1) again gives us the desired recurrence relation for *n*_*k,r*_:

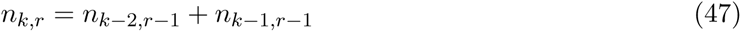

We can now use generating functions (Wilf, 2013) to tackle this recurrence. Define:

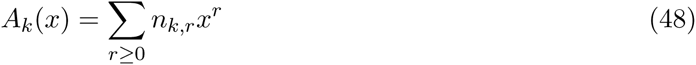
then, multiply both sides of (47) by *x*^*r*^ and sum over *r*. On the LHS:

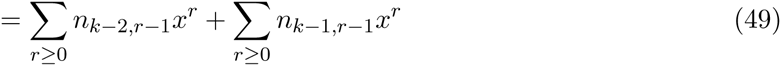

Note that *n*_*k,r*_ = 0 if *r* < 0. Multiplying and dividing the first term by *x* yields:

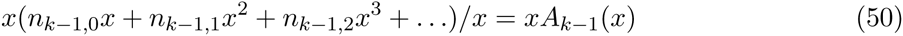

An identical derivation works for the second term. We find:

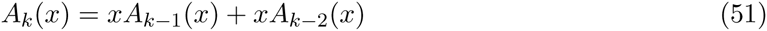

This generating function is in the form of another recurrence.

We can solve this recurrence with the initial conditions below (which can be derived from (47) and its initial conditions):

1. *A*_*0*_(*x*) = 1
2. *A*_*1*_(*x*) = 1 + *x*

The solution to this recurrence with these initial conditions is our desired generating function.

We can see that our generating function works via an expansion and verification the coefficients match known numbers of recombinational meioses for some *k*. For example, let’s expand *g*(*x, k*) at *k* = 5:

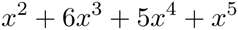
which matches the *n_k,r_* values found via computational calculation.

## C Half-Cousins IBD Length Distribution Simulation Results

Figure 11 show the concordance between our cousin IBD segment length analytic distributions and the binned average (1.98cM bin intervals) of 5,000 simulations.

## D An Approximation of X Pedigree Collapse

Since our models of recent X ancestry omit the possibility of pedigree collapse, it is worthwhile to see when this assumption breaks down. To see how pedigree collapse becomes an increasing problem further generations back, we look at the probability that all of a single individual’s *Ƒ*_*k*+2_ X ancestors, when sampled from a population of *N* individuals with replacement, are distinct. We treat generations as discrete and non-overlapping, and look at the probability that all *Ƒ*_*k*+2_ are distinct individuals as a function of how many generations we go back. This problem is similar to the celebrated birthday problem, but with two rooms of participants: one room of females and another of males. Assuming random mating, each generation, one’s X ancestors must be randomly selected with replacement from a population of *N* individuals. For all ancestors to be distinct, all *Ƒ*_k_ male ancestors selected from a pool of *N*/2 and *Ƒ*_*k*_+1 female ancestors selected from a pool of *N/*2 must be unique:

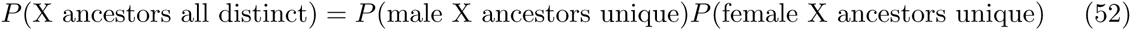

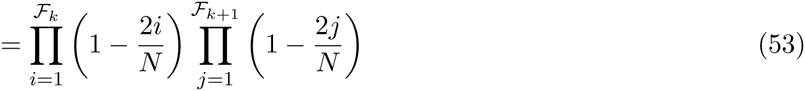

This probability as a function of *k* is plotted in Figure 12. For X ancestors, the probability that at least two individuals are not distinct only becomes a significant problem after around 12 generations. Note that this is a very conservative account of how pedigree collapse could affect our calculations; even if two ancestors were to be non-distinct, this is unlikely to affect our calculations greatly. For pedigree collapse to affect our IBD segment models, an individual has to both be a genealogical ancestor *and* a genetic ancestor of the present-day individual; pedigree collapse has no genetic affect if non-distinct individuals are not genetic ancestors.

For other pedigree collapse related quantities (e.g., what’s the average number of distinct ancestors *k* generations back), see Wachter et al. (1979) approximations which use Feller′s (1950) occupancy models.

**Figure 12:**
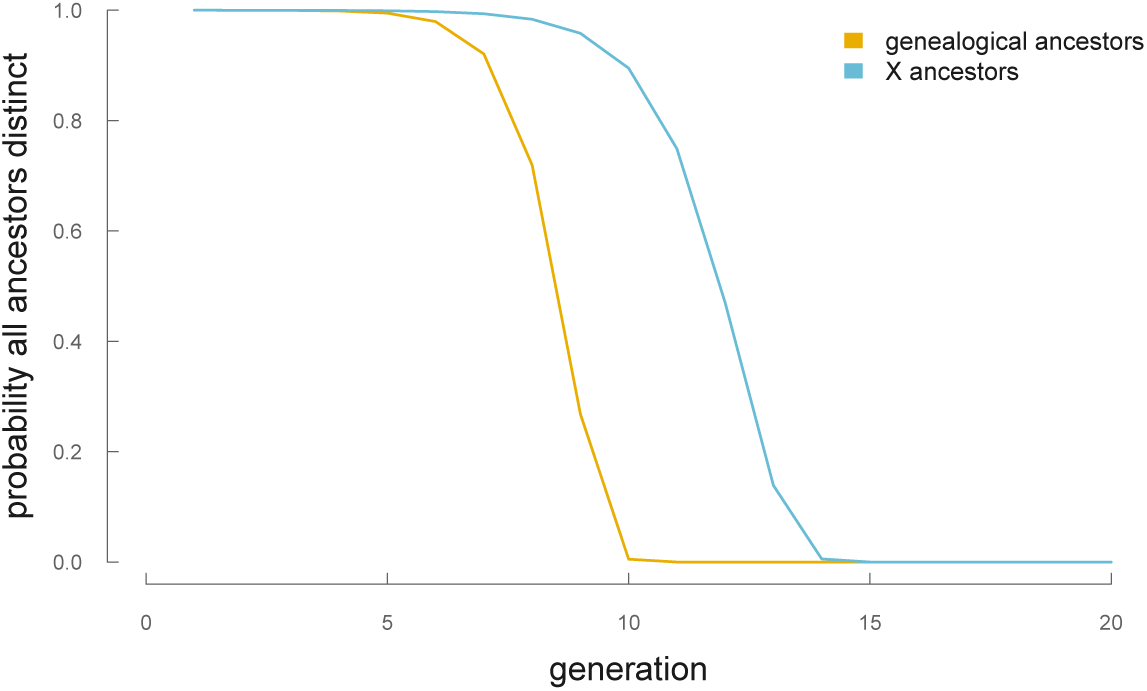
The probability that all genealogical and X ancestors are distinct in a population of *N* = 100,000.

